# Chance and pleiotropy dominate genetic diversity in complex bacterial environments

**DOI:** 10.1101/340828

**Authors:** Lianet Noda-Garcia, Dan Davidi, Elisa Korenblum, Assaf Elazar, Ekaterina Putintseva, Asaph Aharoni, Dan S. Tawfik

## Abstract

How does environmental complexity affect the evolution of single genes? Here, we measured the effects of a set of mutants of *Bacillus subtilis* glutamate dehydrogenase across 19 different environments - from homogenous single cell populations in liquid media to heterogeneous biofilms, plant roots and soil communities. The effects of individual gene mutations on organismal fitness were highly reproducible in liquid cultures. Strikingly, however, 84% of the tested alleles showed opposing fitness effects under different growth conditions (environmental pleiotropy). In biofilms and soil samples, different alleles dominated in parallel replica experiments. Accordingly, we found that in these heterogeneous cell communities the fate of mutations was dictated by a combination of selection and drift. The latter relates to programmed prophage excisions that occurred along biofilm development. Overall, per individual condition, by the combined action of selection, pleiotropy and chance, a wide range of glutamate dehydrogenase mutations persisted and sometimes fixated. However, across longer periods and multiple environments, nearly all this diversity would be lost - indeed, considering all environments and conditions we have tested, wild-type is the fittest allele.

---

The function of most genes may be essential in some conditions, but only marginally contributing or even deleterious in other conditions ^1–4^. The effects of mutations on organismal fitness are therefore environment-dependent, giving rise to complex, pleiotropic genotype-by-environment interactions ^5,6^. (Here, we refer to different effects of the same mutation as ‘environmental pleiotropy’ ^7^, or ‘pleiotropy’ for brevity, and opposing effects in different environments as ‘antagonistic pleiotropy’). Moreover, bacterial populations often do not comprise single cells, but rather have a structure as in biofilms. Under this complexity: changing environments and heterogeneous communities (cell or/and species wise), the fate of mutations could also be dictated by population bottlenecks (drift) or rapid takeover of beneficial mutations in other genes (selective sweeps) ^8–10^. Consequently, the frequency of a given gene allele may change dramatically (from perishing to fixation) with no relation to its protein function ^11,12^.

We aimed at an experimental setup that would examine how complex bacterial growth states and environments might shape protein evolution. Previous systematic mappings were based on a direct linkage between protein stability and function and organismal survival, thus enabling measurements of effects of mutations at the protein level ^5,13–16^. However, how mutations in a single gene-protein affect organismal fitness under varying environments and conditions is largely unexplored ^17^. We thus chose as our model *Bacillus subtilis* NCIB 3610, a non-domesticated strain capable of growing in diverse aquatic and terrestrial environments ^18^. We explored the effects of mutations in different conditions: in dispersed cells in liquid, but also in biofilms where phenotypic and genetic variability prevails ^12^. We also mapped the effects of mutations during spore formation and germination ^19^ and in more complex and close to natural environments including soil, rhizosphere and plant roots.

A catabolic glutamate dehydrogenase (GDH) was our model protein. This enzyme is essential when amino acids such as proline serve as sole carbon-nitrogen sources ^20^. However, in the presence of ammonia and glycolytic sugars, GDH activity is redundant as glutamate must be synthesized rather than catabolized. GDHs therefore respond to changes in carbon-nitrogen sources, and as regulators of glutamate homeostasis, are also associated with biofilm development ^21,22^. *B. subtilis* has two catabolic GDHs, RocG and GudB. The latter is constitutively expressed, and is regulated via association of its hexameric form ^23^. GudB has also regulatory roles ^24,25^ via interactions with the transcriptional activator of glutamate synthase ^25^ and with a transcription termination factor that modulates the stringent response ^26^. We explored mutations in the oligomeric interface of GudB, aiming at multilateral effects on GudB’s enzymatic and regulatory functions.

Altogether, these choices of organism and enzyme allowed us to readily examine and quantify the fate of GudB alleles in a range of different growth conditions and environments, also mimicking natural habitats where strong evolutionary forces act ^12^.

## Experimental setup and data processing

We anticipated that the effects of the explored mutations would be complex and condition-dependent. We thus opted for deep rather than broad coverage and mapped 10 positions within a single ~150 base pairs segment that resides at GudB’s oligomeric interface. The mutagenized positions were arbitrarily chosen while aiming to map highly conserved positions (*e.g.* 58, D in all GudB orthologues) as well as diverged ones (*e.g.* 48, or 61; variable amongst GudB orthologues; **Table S1**). The mutagenized codons were diversified to NNS, whereby N represents any of the 4 bases, and S represents G or C. We thus created 10 libraries, each diversifying a single position into 20 different amino acids *plus* a stop-codon. The libraries were incorporated into the chromosome of *B. subtilis* NCIB 3610 under *gudB*’s original promoter and terminator. The combined library contained 320 single mutant alleles, whose genomes differ, in principle, by a single GudB mutation, including 200 different amino acid alleles (including wild-type), 10 stop-codons, and also synonymous alleles whereby the same amino acid could be encoded by 2 or 3 different codons.

This starting library (the initial mix, hereafter) was used to inoculate cultures grown in an array of different conditions. We tested 7 different growth states with diverse complexity, from single cells to communities: liquid, pellicles (air-liquid biofilms), spores, germinated spores, biofilms grown on agar including on carbon-nitrogen gradients, and colonized soil. Up to 5 different carbon-nitrogen sources were used that, at least as far as the phenotypes of the GudB knockout indicate, inflict different levels of selection on GudB: Glutamate plus ammonia (GA) where ΔGudB has no growth effect; glutamate *plus* glycerol (GG), arginine (A), and arginine *plus* proline (PA), where ΔGudB exhibits a slight growth defect, and proline (P) where ΔGudB exhibits the strongest growth defect (**Fig. S1**). In total, we tested 19 conditions. At each condition, three to five biological replicas were performed by inoculating from the same initial mix. The replicas were grown in parallel, and individually analyzed. Illumina sequencing was applied to determine the frequency of each of the *gudB* alleles in the initial mix and after growth. Following filtering (see Methods), we obtained data for 244 up to 269 individual alleles per experiment (**Data S1** & **Fig. S2**).

The ratio between an allele’s frequency at the end of growth and in the initial mix was derived, and this ratio is referred to as the frequency coefficient (FC; **Data S2**). Basically, FC > 1 means an enriched, beneficial mutation, and FC < 1 a purged deleterious one. However, given the experimental error in determining FC values, values between 0.8 and 1.2 were classified as ‘neutral’, FC ≤0.8 assigned a mutation as ‘deleterious’, and FC >1.2 as ‘beneficial’ (see Methods). Mutations with FC ≤0.1 were classified as ‘highly deleterious’, and similarly, FC ≥10 as ‘highly beneficial’. FC values reflect the relative frequency of alleles, and therefore relate logarithmically to their relative fitness effects (or selection coefficient, *s*). Hence, logFC values were compared throughout. Note, however, that the number of generations differs fundamentally between conditions - *e.g.* ~50 generations in liquid (following 5 serial transfers into a fresh culture) versus effectively no replication in spores (a dormant non-replicative form of *B. subtilis*). Moreover, in pellicles and biofilms, the number of generations cannot be easily determined, and in biofilms different cell types (*i.e.* matrix producers, dormant cells, etc.) have different growth rates ^27,28^. So, while we could not calculate selection coefficients, one should keep in mind that an FC value of 0.8 in spores would effectively mean extinction across 50 generations in liquid (0.8^50^ = 10^−5^).

## Irreproducibility - selection *versus* drift

Our first observations indicated two contrasting scenarios. In liquid cultures, for example, we observed highly reproducible FC values between biological replicas (**Fig. 1a**). Given the small sample numbers (3 replicas as standard, 5 in few cases) the observed variance may underestimate the actual variance. However, the repetitively low variance in a range of different liquid conditions, and in other replica measurements in liquid ^29^, support high reproducibility. In biofilms, however, despite the fact that we did not bottleneck any population upon inoculum, the poor correlation between replicas was evident (**Fig. 1b**). The reproducibility between biological replicas indicates selection, suggesting that in reproducible conditions, the fitness of *GudB*’s and of *B. subtilis* are tightly coupled. In biofilms however, the lack of reproducibility suggested the dominance of drift, i.e., random sampling of GudB alleles.

**Fig. 1.**
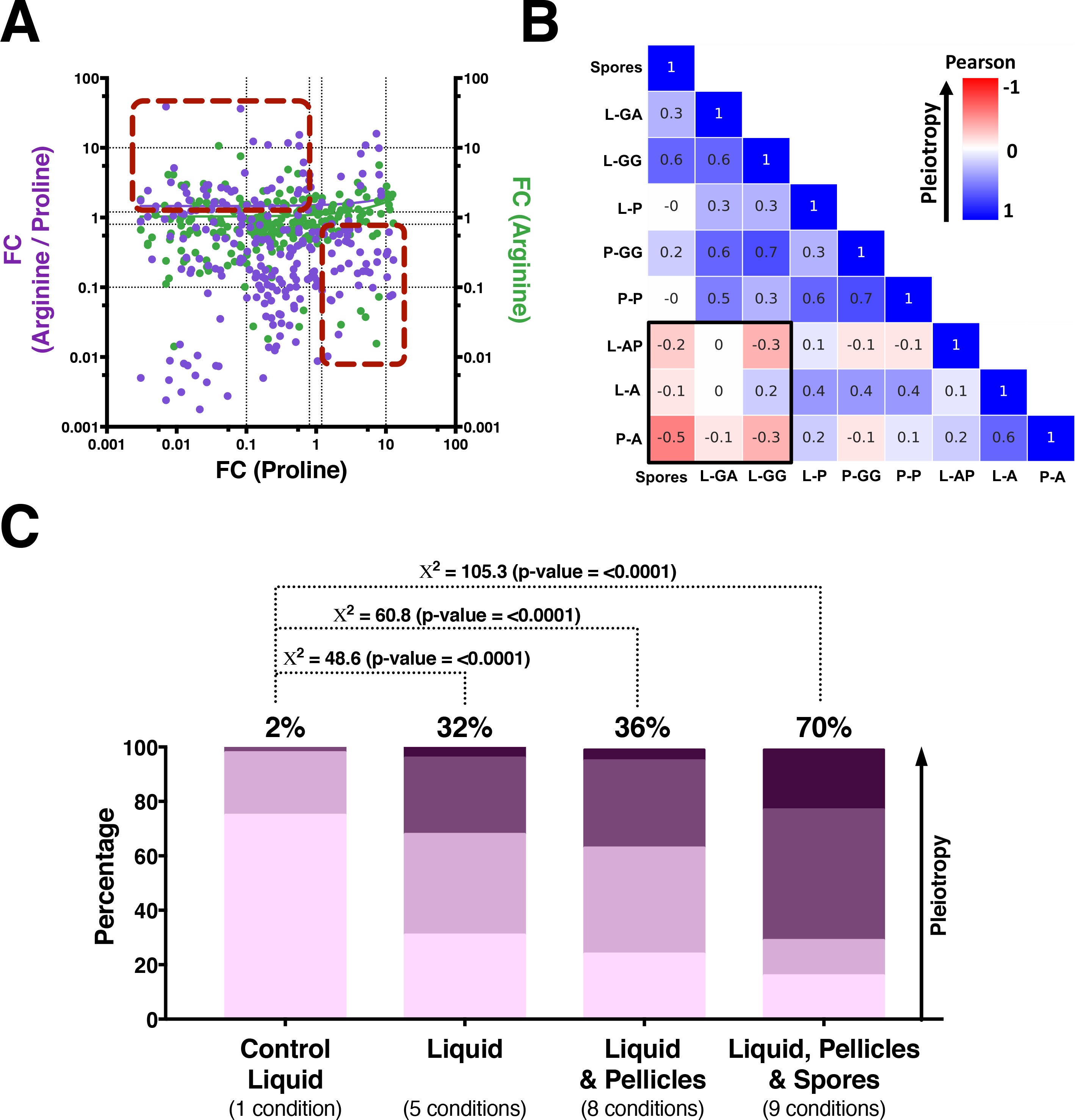
Selection *versus* chance-dominated conditions. **(A)** Dot-plot indicating reproducible measurements of frequency coefficients of individual mutations (FC values) in three parallel replica liquid cultures with proline as carbon-nitrogen source. 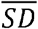 is the average standard deviation between 3 biological replicas. The S.D. values were calculated per each amino acid allele based on logFC values and averaged for all alleles in a given condition. **(B)** The same analysis of three parallel biofilms with arginine as carbon-nitrogen source indicates low reproducibility. **(C)** The 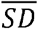 values categorized by the 7 general growth states tested here. Each point represents the 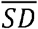 value between 3 replicas of the same experiment (the distributions of SD values per each condition are shown in **Fig. S3a**). **(D)** 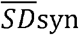 represents the standard deviation between the logFC values of synonymous codons. The standard deviations per allele were averaged for all synonymous alleles in the same replica experiment, and then averaged across the 3 replica experiments in a given condition (the distributions of SD*syn* values per experiment are shown in **Fig. S3b**).

To quantify the contribution of selection versus drift in different conditions, we used two criteria. Firstly, we compared the variability in FC values between replicas by calculating the standard deviation (SD) per allele (using, by default, the logarithm of the FC values; see Methods). The average SD value for all alleles in each experiment 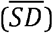 is given for the different growth states (a growth state, *e.g.* liquid, may include several conditions, *e.g.* different carbon-nitrogen sources; **Fig. 1c**; **Fig. S3a** & **Table S2**). As can be seen, in liquid, pellicles and spores, the 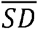 values between biological replicas were low (< 0.06). In biofilms and bulk soil, however, the 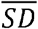 values were > 0.25 indicating low reproducibility. The Fisher test also indicated that for all tested alleles, the variance between FC values significantly changed between liquid, pellicles and spores when compared to germinated spores, biofilms and bulk soil (p values in the range of 0.048 to 1.18 × 10^−27^; **Data S2**).

Secondly, if drift dictates the fate of GudB alleles, codons of the same amino acid should exhibit very different FC values. The deviations between synonymous codon alleles of the same amino acid were calculated, averaged for all alleles in the same experiment, and then for all replicas of the same experiment (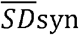, in log values; **Fig. 1d**; **Fig. S3b** & **Table S3**). The Levene’s test confirmed that the variance between synonymous codons is significantly different across conditions (p values in the range of 0.01 to 10^−24^; **Table S4**). Note that the 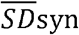 criterion holds within individual replica experiments and is thus independent of the comparison of 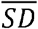 between biological replicas. Nonetheless, these criteria are clearly correlated (**Fig. 1c & d**). Overall, it appears that in liquid, pellicles and spores, the FC values report the outcome of selection acting on GudB alleles at the amino acid level as expected (in few alleles, selection also acted reproducibly at the codon level, **Fig. S4**). In contrast, in biofilms and bulk soil we consistently observed higher 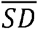 as well as higher 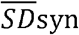 values. In some biofilm experiments, in effect, a single codon had taken over resulting in 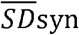 values ≥ 3 (note that logFC values were compared throughout, and the SD for FC values is therefore ≥10^3^).

Given that some conditions were selection-dominated and others were subject to chance, we divided our analysis in two. Firstly, we analyzed selection dominated conditions (liquid, pellicles and spores) to examine whether and how GudB mutations exert different fitness effects under different environments. Secondly, conditions where drift prevailed (germination, biofilms and soil colonization) were analyzed to reveal the relative contributions of selection *versus* chance.

## Pleiotropy - fitness-effects of mutations are condition-dependent

While the FC values, and hence the fitness effects of mutations, were reproducible under many conditions, their distribution varied widely between conditions, including between carbon-nitrogen sources (**Fig. S5**). This indicates pleiotropy - individual GudB alleles have different fitness effects in different environments. To quantify the level of pleiotropy, we compared the FC values of the same GudB mutation across the 9 individual selection-dominated conditions. Because the number of generations differs from one condition to another, we focused on shift from beneficial to deleterious, and vice versa (sign, or antagonistic pleiotropy) because the sign indicates the overall trend irrespective of generation numbers. Representative dot plots comparing the FC values across 3 different liquid conditions are shown (**Fig. 2a**). These indicate that pleiotropy is common, even when comparing liquid cultures with overlapping carbon-nitrogen sources. In particular, a significant number of GudB mutations show antagonistic pleiotropy (dashed squares, **Fig. 2a**). Indeed, the Pearson correlation values for the 36 possible pair-wise comparisons of the 9 reproducible conditions were all below 0.7, and many accommodated a negative value indicating an overall anti-correlation (*i.e.*, dominance of antagonistic pleiotropy; **Fig. 2b**). Across all selection-dominated conditions, 84% of alleles showed antagonistic pleiotropy in at least one of the 36 pair wise comparisons, and 70% of alleles showed mild or strong antagonistic pleiotropy. These pleiotropic effects are far beyond experimental noise, as indicated by comparison to a control sample (**Fig. 2c**).

**Fig. 2.**
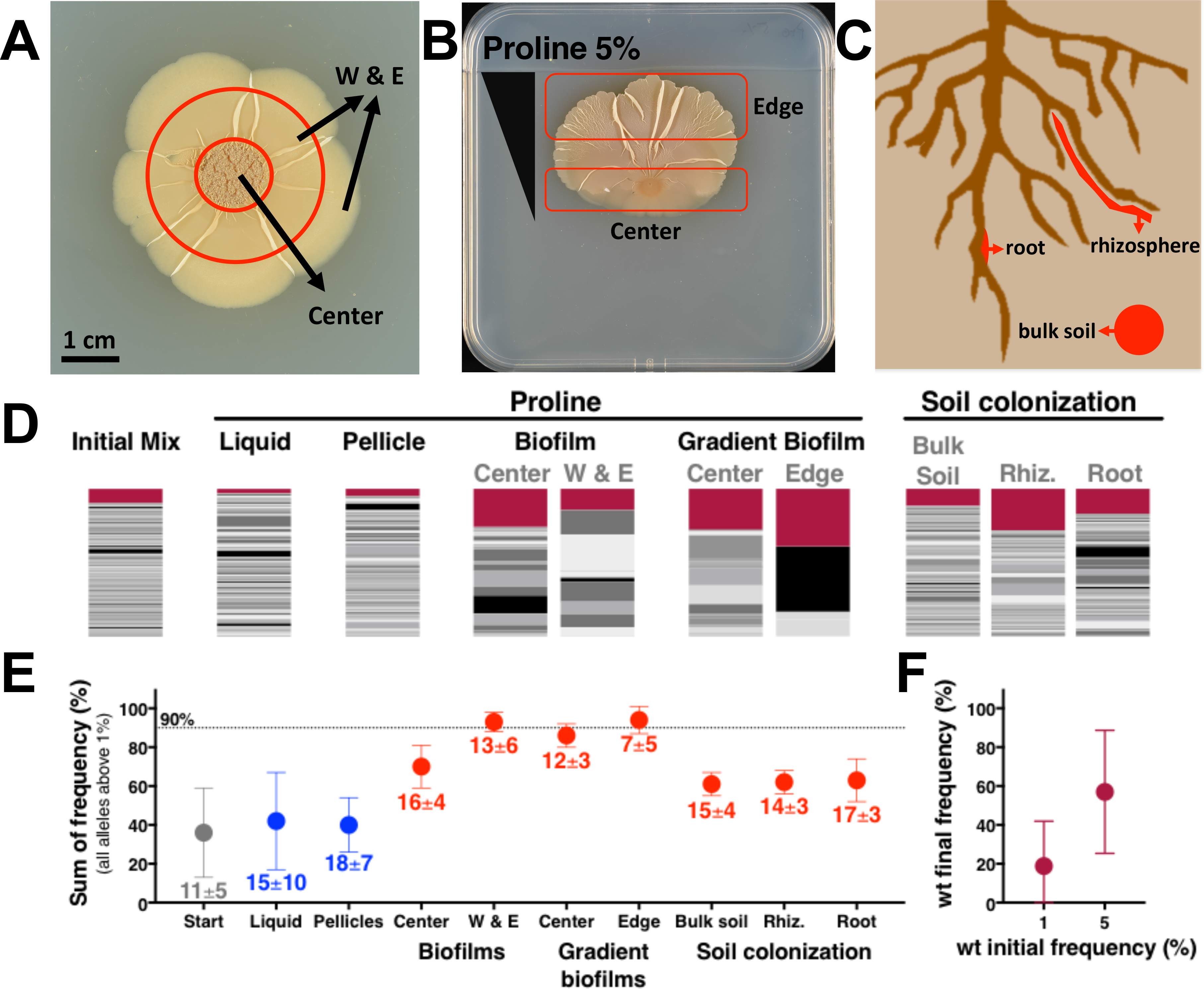
The pleiotropic effects of alleles across different conditions. **(A)** A dot-blot correlation of FC values of individual alleles in three different liquid carbon-nitrogen sources (average values per 3 replicas). The red squares encompass alleles that show sign pleiotropy - i.e., a change from beneficial to deleterious, or vice versa. **(B)** Pairwise correlation of the FC values in all 9 different reproducible conditions (3 replicas per condition; average FC values of all alleles and replicas). Colors indicate the Pearson correlation values (−1, negative correlation; 0, no correlation; 1, positive correlation). The strongest anti-correlation was found with arginine as carbon-nitrogen source (black square). **(C)** The distribution of alleles by their level of sign pleiotropy; from pale to dark purple: (*i*) Weak sign pleiotropy (changes between deleterious and beneficial); (*ii*) Mild sign pleiotropy (changes from highly deleterious to beneficial, or from highly beneficial to deleterious); and (*iii*) Strong sign pleiotropy (changes from highly deleterious to highly beneficial, or vice versa). The fraction of alleles showing mild or strong sign pleiotropy is shown above the bars. The control dataset comprises 4 completely independent growth experiments in liquid proline, each inoculated from a different initial mix and grown on separate occasions (**Fig. S9**). Nonetheless, none of the alleles in this control set exhibited strong pleiotropy. Accordingly, a Chi-squared analysis indicated that the variations between conditions are significantly higher than the variations in the control group (X^2^ and p values are shown^;^ degrees of freedom equal 3 in all cases).

Overall, the dominance of pleiotropy meant that across all conditions where selection acts, 86% of the alleles were beneficial in at least one condition. However, not a single mutation was beneficial across all conditions. Further, if a mutation were to be considered deleterious if purged under at least one condition, then 98% of the tested GudB mutations were deleterious.

## Combined action of selection and drift in heterogeneous environments

In biofilms (and also in germination and soil colonization though to a lesser degree), irreproducibility between replicas, variability between codons (**Fig. 1c & d**), and the near-fixation of relatively few alleles (**Fig. 3**), all suggested fixation by chance. What is the nature of these few GudB ‘winners’, are they merely lucky?

**Fig. 3.**
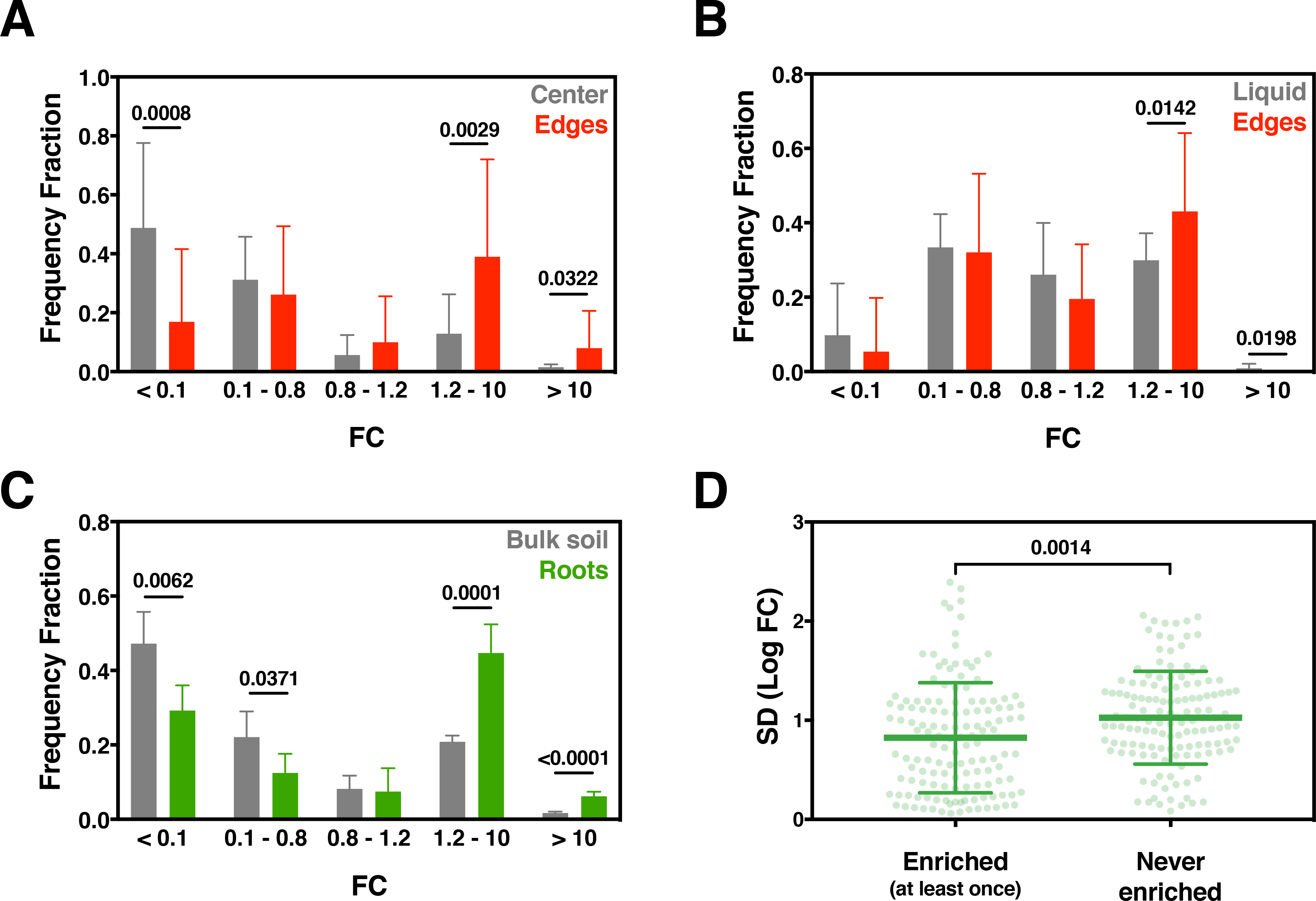
Genetic sweeps in biofilms and soil and the dominance of the wild-type allele. Photographs of 5 days old biofilms: (**A**) normal biofilms; (**B**) gradient biofilms; (**C**) a scheme of soil colonization (shown biofilms with proline as carbon-nitrogen source). (**D**) The distribution of frequency of individual alleles for different growth states with proline as carbon-nitrogen source. Bar widths represent allele frequency from raw read counts (Rf values; **Data S1**). Magenta corresponds to wild type GudB. (**E**) Alleles with Rf ≥ 1% were identified and their number and sum of frequencies are shown (averages and standard deviations for all experiments in a given condition). Blue designates selection-dominated conditions and red drift-dominated ones, as in **Fig. 1**. (**F**) An initial mix of 4 alleles was created including the wild-type allele at a varying frequency from 1 up to 25%. Following growth in normal and gradient biofilm with proline, the frequency of wild-type reached an average of 18% when initiated at 1%, and up to 100% when initiated at 5% (see **Fig. S10** for the entire dataset).

While drift dominated in biofilms and soil colonization, curiously, wild type GudB was enriched in up to 85% of these experiments suggesting that selection may also play a role (**Fig. 3**). To assess the action of selection, we compared the three biofilm areas. There appears a systematic trend, whereby alleles enriched in the edge are more likely to arise from alleles that persisted or even enriched in the center (**Fig. 4a**). Similarly, 75% of the enriched edge alleles were neutral or beneficial under liquid growth with proline, a condition under which GudB experiences the strongest selection (**Fig. 4b**). This suggested that although drift dominated GudB’s fate in biofilms, GudB was under selection at some stage of biofilm development. Accordingly, we found that in biofilm centers the FC values are less skewed and more reproducible than in the edge or wrinkles (**Fig. S5b**) and the center 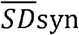 values are half (**Table S3** & **Fig. S3b**). The 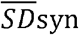 values are obviously much higher in the biofilms’ center compared to liquid cultures, but the trend suggests that at the onset of the biofilm’s development, selection acts on GudB (**Table S3** & **Fig. S3b**). Foremost, that wild type GudB was present at high frequency in the vast majority of biofilm experiments is not a coincidence that relates only to its high frequency in the initial mix. Indeed, a spiking experiment indicated that wild type was enriched by nearly 20-fold even when scarcely present in the initial mix (**Fig. 3f**).

**Fig. 4.**
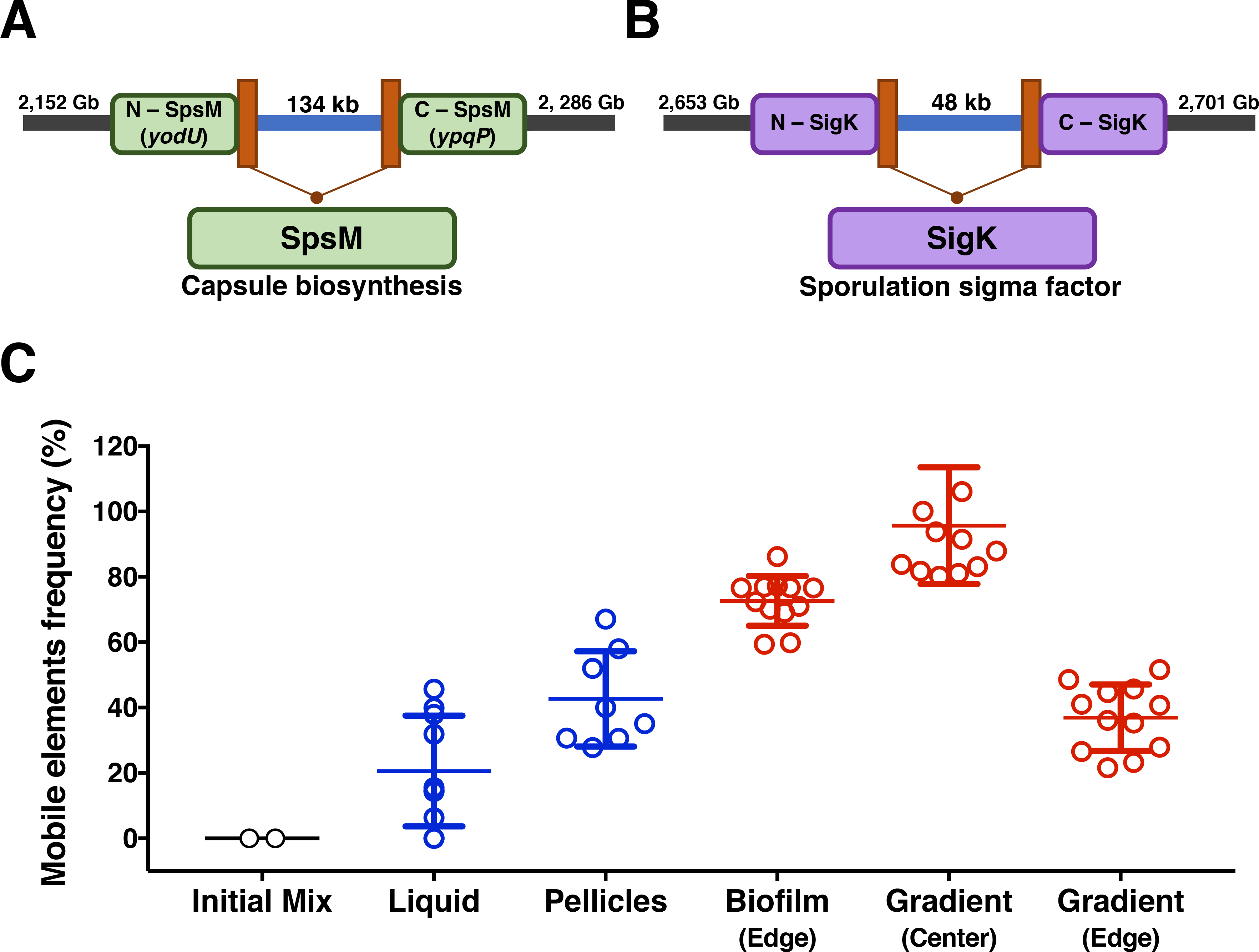
The combined action of selection and chance in biofilms (red) and soil colonization (green). **(A)** Alleles that enriched in the edge of the biofilms are more likely to arise from alleles that were neutral or enriched in the center. The distribution of categorized FC values from all biofilm centers (grey) compared to the distribution of FC values of center alleles that were enriched in the edge (red). **(B)** The distribution of categorized FC values of all alleles in all liquid conditions (grey) compared to the distribution of FC values of liquid alleles that were enriched in the edge of biofilms (red). **(C)** Alleles enriched in the root are more likely to arise from alleles that were enriched in the bulk soil. The distribution of categorized allele FC values in all soil samples (grey) compared to the distribution of FC values of soil alleles that were enriched in the root (green). **(D)** The distribution of SD values (variability between replica experiments, as in **Fig. 1c**) of alleles enriched in one or more root populations compared to alleles that were never enriched in the roots. T-tests were computed per each FC category. p-values indicating significance (p < 0.05) are presented above the bars (details of all the T-tests are provided in **Table S7**).

Similarly, we searched signatures of selection in soil colonization - a process that involves multiple passages, beginning with a change of medium (Hoagland solution rinses; see Methods) and results in colonization of the roots. As in biofilms, there is a statistically significant trend whereby alleles enriched in the root are more likely to arise from alleles that were enriched in the soil (**Fig. 4c**). Further, 19 amino acid alleles were found to be enriched in at least 10 out of the 15 sequenced populations, suggesting some degree of reproducibility (**Table S5**). Selection during soil colonization is also manifested in the variation between biological replicas (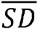 values) of alleles that were enriched in root populations being on average 20% smaller than those that were not (**Fig. 4d**). Finally, stop codons were purged in all biofilms and soil populations, indicating, as expected, that GudB’s activity is required for *B. subtilis*’ survival under these conditions ^22,23^.

Altogether, in biofilms in soil colonization both drift and selection determine the fate of GudB alleles. We further examined the biofilms as described in the next section.

## Drift in biofilms relates to programmed prophage excisions

Mutagenic rates in biofilms are high and mutations with a selective advantage rapidly take over (genetic sweeps) ^30,31^. Growth in biofilms is also spatially defined, giving rise to segregated lineages whereby an entire segment of the biofilm’s edge stems from a single cell in which a beneficial mutation had first emerged ^12^. GudB mutations that happen to be in these ‘founder’ cells might therefore fixate along these lineages. In pellicles that are largely considered as biofilms, intercellular matrix is produced ^32^ but spatial segregation is less pronounced than in solid biofilms. Accordingly, we found that in oppose to agar biofilms, in pellicles selection acts reproducibly (**Fig. 1**). To further establish that spatial segregation is a key factor, we divided the edges of the biofilm into small sections, and sequenced them. We found that most sections contained a single GudB allele (**Fig. S6**). Thus, in a way, the GudB allele represents a ‘barcode’ that reports single founder cells giving rise to individual sectors of the biofilm ^12^.

What might be the mutations driving these genetic sweeps and spatial segregation? We sequenced samples for which enough genomic DNA was available (6 ordinary and 12 gradient biofilms, and for comparison, 2 initial mix, 6 liquid and 4 pellicle samples). A range of single nucleotide polymorphisms (SNPs) in various loci was identified across these samples (**Data S3**). We focused, however, on identifying genomic mutations that were not, or scarcely observed in the initial mix and/or in liquid samples, suggesting that they emerged and enriched in the biofilms.

Foremost, we observed two large genome deletions that occurred in all biofilms with a frequency approaching 100% (**Fig. 5a** & **5b**). These deletions correspond to the excision of two mobile genetic elements, or prophages, **skin** and SP-β ^33–35^. Excision of *skin* generates a functional protein: sigK - a sporulation-specific transcription factor essential for cell differentiation in *B. subtilis* ^36^. The excision of SP-β generates another functional protein dubbed SpsM - a protein involved in capsid polysaccharide biosynthesis mediating ^37^ and with relevance to pellicle development ^38^. Nearly all biofilm cells carried one of these variations, and most cells carried both (**Fig. 5a** & **b; Table S6 & Data S3**). These prophage excisions therefore appear to be under stronger selection than the GudB mutations. The frequency of these structural variations gradually increases, from none in the initial mix to 100% in gradient biofilms (**Fig. 5c**), and so does the signature of GudB’s drift (**Fig. 1**). However, at this stage, the observed link between the prophage excisions and GudB’s drift is circumstantial and further experiments are needed to establish how are these two phenomena linked. The prophage excisions are also likely to occur in the soil, but the DNA recovered from these samples was insufficient to allow genome sequencing.

**Fig. 5.**
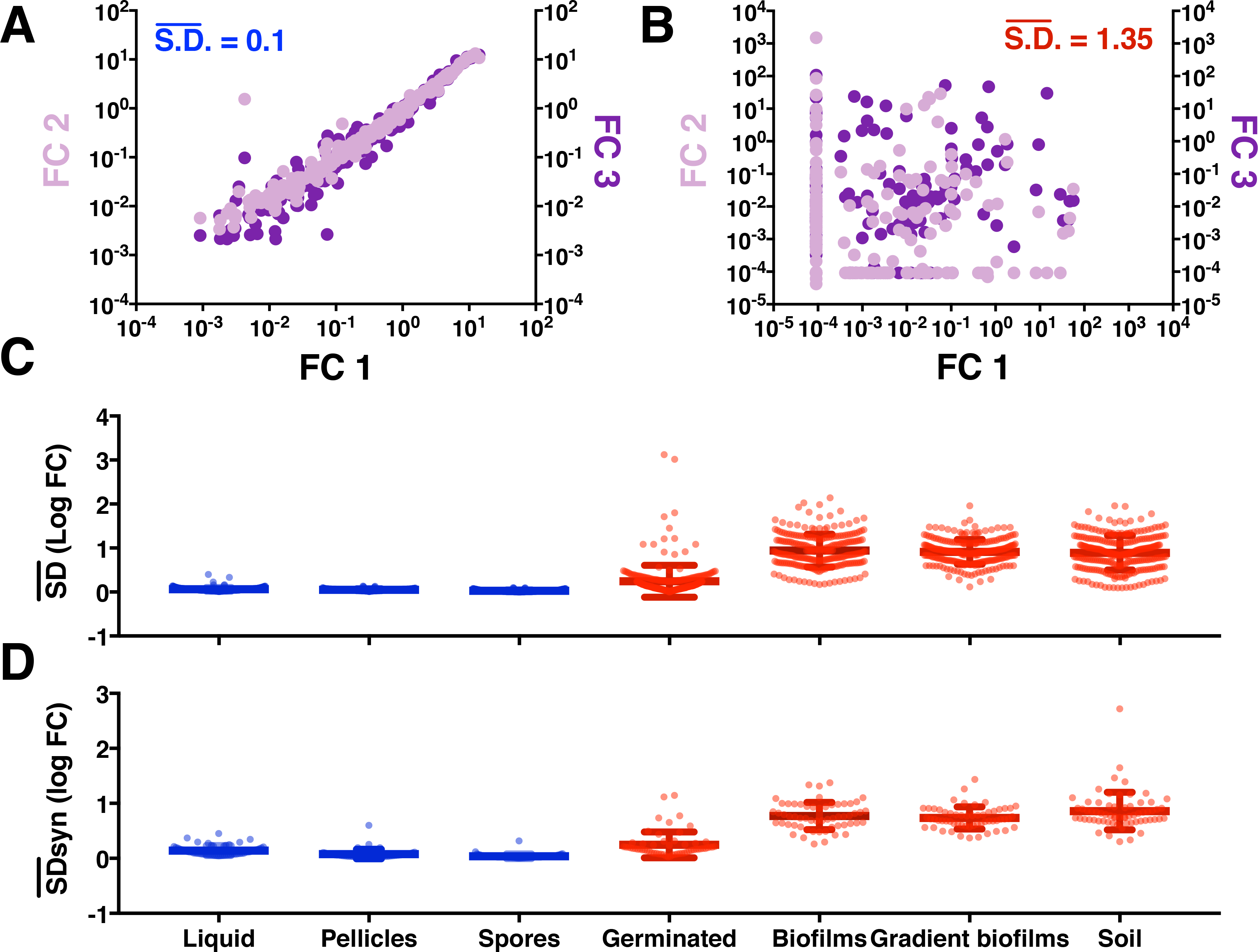
Programmed genomic excisions drive GudB’s drift in biofilms. Schematic representation of *B. subtilis* genomic organization before and after the excision of the prophage mobile elements SP-β (**A**) and skin (**B**) and their position in the genome. (**C**) These excisions were absent in the initial mix yet dominated biofilms and went to near fixation in the edge of gradient biofilms (for frequencies in individual experiments see **Table S6**). Excision of the mobile elements occurred in two different genomic locations within the same experiment. The values were summed and averaged according to the general condition shown. The details of the excisions (location and frequency) per experiment are shown in **Data S3**.

Exclusively in biofilms, we also detected 59 enriched SNPs in a conserved region of 16S rRNA (**Table S6** & **Data S3**). However, *B. subtilis* has ten 16S rRNA gene copies. Since these are essentially identical, we could not determine which of these 10 paralogues carried mutations. However, per population, 98% of the 16S rRNA mutations occurred in the same Illumina read suggesting that one paralogue was highly mutated while others remained intact (**Fig. S7**). Large differences in expression levels of 16S rRNA genes were identified **in P. aeruginosa** biofilms ^39^, and ribosomal heterogeneity has been linked to biofilm development in *B. subtilis* ^40^. Yet, to our knowledge, mutations in the 16S rRNA genes have not been reported in biofilms. At this stage, however, which 16S gene is inactivated, how do multiple proximal 16S mutations occur, and how inactivation affects biofilm development, remains unclear. Overall, the 16S rRNA SNPs, and the structural variations in particular, seem to have a key role in biofilm development in *B. subtilis*. Accordingly, most of these genetic variations were reproducible between replica experiments (**Table S6**) suggesting that they arose during biofilm growth and then enriched by virtue of promoting biofilm formation ^12^.

## Concluding remarks

That the fitness effect of a mutation may vary depending on the environment is generally assumed. However, the magnitude of environmental pleiotropy unraveled here is surprisingly high. Environmental changes, including minute ones like addition of arginine to a proline medium, can completely revert the effect of *GudB* mutations. Overall 84% of the tested GudB mutations showed sign reversions. Pleiotropy severely restricts protein sequence space. Extensive pleiotropy has an interesting implication. The so-called wild type sequence of a gene-protein is generally thought to represent just one sequence out of an entire cloud of related sequences that are similarly fit. However, our results indicate that wild-type GudB’s sequence is singular in being fit across multiple constrains and environments. Per individual tested conditions, most mutations are either neutral or beneficial (28 - 81%). However, if all tested conditions are considered, only 2% of the tested GudB mutations are neutral or beneficial in the 9 reproducible conditions when selection acts on GudB.

The pleiotropy of protein mutations across multiple growth and environmental conditions has been rarely measured ^17^, and, to our knowledge, never for a protein with an intrinsic physiological role. Indeed, the extensive pleiotropy observed here might be the norm in cases of complex relationships between a protein’s expression and activity levels (‘protein fitness’) and organismal fitness, as with GudB. The high degree of pleiotropy observed here may also relate to GudB’s role as an enzyme and regulator, and also to the positions explored (oligomer interface). In any case, our results suggest that, as currently performed, laboratory mutational scans broadly underestimate the fraction of mutations that are deleterious in ‘real life’.

Together, pleiotropy and drift dictate not only the evolution of short-term polymorphism (micro-evolution) but also the evolution of protein sequences along long evolutionary times and across species (macro-evolution). Indeed, the correlation between the effects of mutations in laboratory mappings and their occurrence or absence in natural sequences is limited ^13^. However, laboratory mappings represent a single condition, and merging of data from multiple conditions could in principle reveal higher correlation. Identifying trends in complex datasets requires an unbiased approach. However, even when several different machine learning approaches were applied, merging of conditions gave no further correlation (**Fig. S8**). Thus, along short evolutionary periods, proteins experience variable and opposing selection pressures. Additionally, drift may lead to rapid fixation of alleles that are marginally fit or even deleterious. The effects of drift have been extensively studied initiated by Kimura’s neutral theory ^11^. Our results quantify its effect in bacterial populations and the potential effect of drift in combination with selection across different environments. For example, nearly 80% of the tested mutations survived or even enriched during sporulation, and a single spore could then initiate a whole new population. However, once the environment changes, such alleles will be rapidly lost unless compensated or a priori enabled by other mutations. Compensation, or enabling by other mutations, results in epistasis, i.e., in the effect of mutations being dependent on the sequence context in which they occur ^41^. Accordingly, along macro-evolutionary time scales, epistasis dominates gene and genome sequences ^42^.

## Acknowledgments

L.N.G. was supported by the CONACYT grant #203740 and the Martin Kushner Fellowship at the Weizmann Institute of Science. D.S.T. is the Nella and Leon Benoziyo Professor of Biochemistry. Financial support by the Kahn Center for Systems Biology at the Weizmann Institute of Science is gratefully acknowledged. We are highly grateful to Ron Milo, Sarel Fleishman, Zvi Livneh and Fyodor Kondrashov for support and critical advice, to Einat Segev, Arjan de Visser for critical and insightful comments to the manuscript. We highly appreciate the help of Moshe Hershko in script development for data processing, and of Yinon Bar-On, Shmuel Gleizer in the analysis of genomic sequences. We are grateful to Ron Rotkopf from the Weizmann Life Sciences Core Facilities for his guidance on the statistical analysis. We are thankful for the services provided by the Crown Genomics institute of the Nancy and Stephen Grand Israel National Center for Personalized Medicine, Weizmann Institute of Science.

## Authors Contribution

L.N.G. and D.S.T. designed experiments and wrote the manuscript. L.N.G., D.D. and D.S.T. analysed the data. L.N.G. performed all experiments, except selection in soil colonization that was performed in collaboration with E.K. and A.A. D.D. and A.E. wrote the scripts used for data analysis and visualization. E.P. applied machine learning classification.

## Materials and Methods

### Strains

*B. subtilis* NCIB 3610 DS7187 (kindly gifted by Dr. Daniel B. Kearns ^43^) that lacks the ComI peptide and has high competence capacity similar to domesticated *B. subtilis* strains was recruited to this study. *Bacillus subtilis* NCIB 3610 *gudB::tet* strain ^23^ genomic DNA was transformed into *B. subtilis* NCIB 3610 DS7187. *B. subtilis* NCIB 3610 *ΔcomI gudB::tet* was thus isolated, and was phenotypically and genetically tested.

### GudB allele library construction

We performed site directed mutagenesis in 10 codons (amino acids: M46, L48, K52, D58, D59, S61, K63, T66, Y68, S75) of the *gudB* gene cloned in the pDG_GudB plasmid, which was modified from the pDG1728 backbone vector ^23^. The codons were mutated to NNS (N = all bases & S = C or G) whereby the 20 standard amino acids and 1 stop codon is encoded. The codon mutagenesis was done in one step PCR protocol and independently for each position. Thus, we created 10 libraries, each containing 20 different amino acid alleles (non-synonymous, missense mutations), 1 stop-codon (nonsense), and 11 synonymous alleles (alternative codons encoding the same amino acid). All mutagenic PCRs were performed with Kapa HiFi HotStart Ready Mix (Kapa Biosystems) following manufacturers conditions (**Table S8** shows the sequence of all primers). The 10 PCR products were purified and used to transform the *E. coli* T10 strain (Thermo Fisher Scientific). Clones were pulled together after an overnight growth on LB + Ampicillin (100 μg/ml) agar plates at 37°C. At this stage, 4 to 6 clones per library were isolated and analyzed by sequencing. Total plasmid DNA from these library transformations was extracted and also analyzed by sequencing. Each of the 10 libraries contained, after transformation, at least 10^5^ clones, corresponding to ≥ 1000-fold coverage per allele. Approximately 10 μg of plasmid DNA, from each library, was linearized (XhoI, New England Biolabs, following manufactures conditions), purified, and used to transform the *B. subtilis* NCIB 3610 *gudB::tet ΔcomI* strain. Transformations were performed as described ^23^. After transformation, overnight growth on in + Spectinomycin (100μg/ml) + Glucose (0.5 mg/ml) agar plates was used as selection. The resulting cells were pulled together and kept at −20°C in 50% glycerol. In total, 10 *B. subtilis* libraries were constructed in parallel and each contained, after transformation, at least 10^4^ clones (≥100-fold coverage per allele). Genomic DNA extraction of each library was performed (GenElute - Sigma). The integrity of the mutagenic process was verified by sanger sequencing the *amyE::gudB* locus indicating that mutations were observed only in the diversified codon.

### Selection and growth conditions

10 ml of LB (1% tryptone, 5% yeast extract and 1% NaCl) with Glucose (0.5%), ammonium sulfate (0.5%) and spectinomycin (100 μg/ml) cultures were inoculated with 1 ml of each library stock. The cultures were grown overnight at 37°C with shaking. 500 μl of the overnight culture was used to inoculate 3 ml of LB plus glucose (0.5%) and ammonium sulfate (0.5%). The cultures were incubated at 37°C with shaking and once the O.D._600_ reached 0.8 they were mixed equally and used as the starting population (initial mix). A fraction of the cells at this stage were harvested by centrifugation and stored for genomic DNA purification. In total, three different initial mixes were used for the experiments described here. Initial Mix #1 was used to inoculate most liquid conditions (4 carbon-nitrogen sources), pellicles and gradient biofilms. Initial Mix #2 was used to inoculate 1 liquid condition, spores, germination and biofilms, and initial mix #3 was used to inoculate bulk soil (**Data S1**). Detailed selection conditions are listed below:

For selection under liquid serial passages 100 ul of the initial mix was used to inoculate 10 ml cultures of MS medium (5 mM potassium phosphate, 100 mM MOPS pH 7.1, 2 mM MgCl_2_, 700 μM CaCl_2_, 50 μM MnCl_2_, 50 μM FeCl_3_, 1 μM ZnCl_2_, 2 μM thiamine, 50 μg/ml tryptophan, 50 μg/ml phenylalanine and 50 μg/ml threonine)^23^ with glucose (0.5%) plus ammonium sulfate (0.5%), glutamate (0.5%) plus glycerol (0.5%), proline (0.5%), arginine (0.5%) or proline (0.25%) plus arginine (0.25%). The cultures were incubated at 30°C with shaking until O.D._600_ reached 1 - 1.5, after which 100 μl was used to inoculate 10 mL of fresh medium. The serial passages were done every 24 hours when proline (0.5%), arginine (0.5%) or proline (0.25%) plus arginine (0.25%) where used as carbon-nitrogen sources, and every 12 hours when glucose (0.5%) plus ammonium sulfate (0.5%), or Glutamate (0.5%) plus glycerol (0.5%), were applied. In total, all liquid passages were maintained for approximately 50 generations.

For selection in pellicles, 100 ml of media (b), (c) and (d) were inoculated with 100 μl of the initial mix cells. The culture was incubated at 30°C without shaking, for 5 days.

For selection of spores and germinated spores, three ml of the initial mix was used to inoculate 25 ml of Difco Sporulation Medium (DSM) in 250 ml flasks and incubated at 37°C with 150 rpm shaking until O.D.600 reached 0.4. This culture was used to inoculate 250 ml of fresh DSM in 1L flasks. The cultures were incubated 48h at 37°C with 150 rpm shaking. Cells were subsequently harvested by centrifugation and stored at 4°C over night. After, cells were re-suspended with 200 ml of cold deionized sterile water (dW) and incubated for 30 min at 4°C. Cells were harvested and re-suspended with 200 ml of cold distilled water (dW) and incubated overnight at 4°C with slow orbital agitation, to kill all planktonic of vegetative cells. The culture was harvested, re-suspended in 30 ml of dW and heated to 80°C for 20 min. Finally, spores were harvested, re-suspended in 10 ml of dW, and stored at −20°C. To germinate these spores, they were diluted 1000 times in phosphate-buffered saline solution and 100 μl of this suspension was used to inoculate LB plus glucose (0.5%) agar plates (10 plates). Approximately 10,000 colonies were obtained and pulled together.

For selection in biofilms, MS agar (1.5%) plates supplemented with different carbon-nitrogen sources were prepared. For gradient biofilms, gradient agar plates were prepared. First, square plates (12×12 cm) with MS agar (1.5%) medium were poured. After the agar solidified, an area of 2×14 cm was removed from the top of the plate. In this area, a solution of either Proline 5%, Arginine 5%, monosodium glutamate 5% or Glycerol 5% in 1.5% agar was poured into the removed section. For the glutamate plus glycerol gradient biofilm, two opposite areas of the agar plate were removed. Into one, a solution of monosodium glutamate (5%) in 1.5% agar was poured, and into the other, glycerol (5%) plus 1.5% agar solution (see **Fig. S11a** for a graphic representation of the agar plates preparation). All gradient agar plates were incubated for 24 h at room temperature before use. We also calibrated the place in the gradient plate where we inoculated the cells such that we observed growth after 1 night incubation at 30°C (**Fig. S11b**). For growth in biofilms and gradient biofilms, 5 μl of the initial mix were used as inoculum. Plates were incubated for 4 days at 30°C and 2 more days at room temperature. The colony was then dissected in 3 areas (center, wrinkle and edge) for normal biofilms, and in 2 areas (center and upper) for gradient biofilms (illustrated in **Fig. S11c-g**). After selection in all the above-mentioned conditions the biomass was harvested and storage at −20°C. All growth experiments were performed in triplicate by inoculating with the same initial mix.

For selection in soil and plant roots, the initial mix was generated as above-mentioned except that the process was scaled up (instead of 3 ml, 10 ml of culture was prepared per library). In total, 200 ml of the initial mix (O.D._600_ = 0.8) was applied. This LB culture was washed three times (by means of centrifugation and re-suspension) with 100 ml half strength Hoagland solution ^44^. After the final wash, the cells were re-suspended in half strength Hoagland solution to a final O.D._600_ of 0.1. Since Hoagland’s solution is not isotonic, the washes resulted in death of about a third of the *B. subtilis* cells. Thus, handling the samples at this stage was performed as fast as possible. The Hoagland solution imposes some selection pressure on the initial mix population although the loss in population size is relatively small (≤30%). The soil colonization FC values therefore result from the entire process that begins the rinses with the Hoagland solution and ends with the colonization of the roots. Natural soil was collected at the Ha-Masrek Reserve, Israel (31.793 N, 35.042 E), sifted through 2 mm sieve and autoclaved three times for 30 min at 121°C. A total of five pots (size 10 × 8 × 5 cm) with autoclaved natural soil were drenched with the initial mix suspended in half strength Hoagland Solution ^44^. These potted soils drenched with bacterial suspensions were used to plant tomato seedlings grown first in sterile conditions. Seeds of tomato (*Solanum lycopersicum L*.; cv. Micro-Tom) were surface-sterilized with 70% ethanol for 5 minutes and, 10 minutes with 3% bleach with 0.01% Tween 20. Surface-sterile seeds were germinated on sterile filter paper (Whatman, catalog # 1001-085) saturated with half strength Hoagland Solution for 7 days (23°C and 16 hours photoperiod). Six tomato seedlings were transferred to each pot and grown for one month (21°C, 16h light, 8h dark) with drenching with half strength Hoagland twice a week. Plants were subsequently harvested from the five pots. Roots and rhizosphere samples were collected for each replica experiment consisting a pool of six roots. First, the plants were carefully removed from the soil. Roots were then cut out from the plants and vortexed in 20 ml of washing solution (0.85% NaCl) for 30 s. This step was repeated one more time with a fresh washing solution. The combined root washing solutions (40 ml) was centrifuged for 30 min at 3000 rpm and the resulted pelleted samples corresponding to the rhizosphere were frozen in liquid nitrogen and stored at −80°C. The washed roots were blotted in filter paper and stored at −80°C until further use. Finally, bulk soil without roots was also stored at −80°C.

### Genomic DNA extraction

All samples, including pellicle, spores, biofilm and gradient biofilm samples, were defrosted and re-suspended in 10 ml of dW. The samples were sonicated at 40% power, VibraCell, Sonics, for 10 min at 60 s intervals. Cells debris was harvested by centrifugation (13,000 g for 20 min). Genomic DNA from all samples was extracted using the GenElute Bacterial Genomic DNA Kit (Sigma-Aldrich) generally following the manufacturer’s instructions, with the exception of the soil, rhizosphere soil and plant roots samples. For these samples, the PowerSoil DNA Isolation kit of Mo Bio was used, following its manufacturer’s instructions.

### Illumina sample preparations

The mutagenized *gudB* fragment (from amino acids 45 to 81) was amplified using the primers GudB_In_For (5’-CTCTTTCCCTACACGACGCTCTTCCGATCTnnnnnnCCCGAAGAGGTATACGAATTGTTAAAAGAG), and GudB_In_Rev (5’-CTGGAGTTCAGACGTGTGCTCTTCCGATCTCGCCTTTCGTTGGACCGAC). To the GudB_In_For primer, 6 N’s were added to increase the sequence variability between amplicons. PCRs were performed with the KapaHiFi HotStart Ready Mix (Kapa Biosystems) using approximately 100 ng of genomic DNA as template and following manufacturer’s instructions. Using 10 μl of the PCR as template, a second PCR was performed to add the Illumina adaptor sequence, using primers GudB_Out_For (5’- AATGATACGGCGACCACCGAGATCTACACTCTTTCCCTACACGACGC) and GudB_Out_Rev (5’- CAAGCAGAAGACGGCATACGAGAT<u>TGTTATAC</u>GTGACTGGAGTTCAGACGTGTGC). The Illumina index (underlined) was changed in the GudB_Out_Rev primer to different Illumina indexes. Each condition was differently barcoded. All PCRs were purified using the Agencourt AMPure XP (Beckman Coulter). The concentration of PCR products was verified using Qu-bit assay (Life Technologies).

### Analysis of the Illumina reads

DNA samples were run using the Illumina NextSeq 150-bp paired-end kit. The FASTQ sequence files were obtained for each run and customized using MatLab 8.0 and Python 3.6 scripts designed to count the number of each individual allele in each sequenced sample. We filtered the reads to exclude any reads that have mutations outside the mutagenized codons. All codons encoding for the wild-type amino acid were summed in one and assigned as WT. All other codons were counted independently. The unprocessed read counts are shown in **Data S1**. Further filtering excluded alleles with < 100 counts in the initial mix to avoid statistical uncertainty with respect to FC values. In total, we obtained data for up to 269 individual alleles per condition out of the originally introduced 320 alleles. Per condition, a minimum of 380,000 reads was obtained. Thus, in average, we obtained 1500 reads per allele.

### Data Analysis

The frequency of each allele (f_i_) was calculated as the ratio between the number of reads for allele i divided by the total number of reads. The allele frequency coefficient (FC_i_) was subsequently calculated as the ratio of after selection (f_i_) divided by the frequency of the same allele in the initial mix (**Fig. S2** & **Data S2**). Normalization by the number of wild-type reads rather than by the total number of reads gave essentially identical FC values for the majority of samples. However, in the few samples where wild-type frequency was significantly reduced after selection, normalization resulted in high noise and large biases including large changes in sign (higher sign pleiotropy). FC values were therefore derived from the unnormalized frequency (fraction of reads for a given allele out of the total number of reads). FC values relate to fitness logarithmically, and thus logFC values were compared. To this end, all FC’s equal to zero had to be changed, and we opted for a tenth of the minimum FC value found amongst all experiments. For the liquid, pellicles, biofilms, spores and germinated spores experiments (**Data S2, sheet 1**) the zeros were changed to 4.2 × 10^−6^. For the bulk soil experiments (**Data S2, sheet 2**) zeros were changed to 1.14 × 10^−5^. The logarithm of all FC values was calculated and was also used to derive mean FC values. The logFC values were then used to calculate: (i) the standard deviation for all alleles across conditions (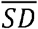; the standard deviation between logFC values observed per each allele in replica experiments were averaged for all alleles measured in a given condition); (ii) the standard deviation between synonymous codons within the same replica experiment (deviations between logFC values of synonymous codons of the same amino acid allele were calculated, averaged for all alleles in the same experiment, and then for all replica experiments per condition). The T, F, X^2^ tests and Pearson correlation values were obtained using the PRISM software. The Levene’s test was performed using R. In addition, **Table S9** from supplementary material shows the sample size for every test performed in this study.

### Defining the limits of neutrality

From all conditions tested here, only in glucose plus ammonia the GudB knockout had no growth effect (**Fig. S1**). Hence, this condition is largely neutral, and the variation observed in FC values would primarily be the outcome of noise. The standard deviation between 3 biological replicas was calculated per allele, and these values spanned over the range of 0.002 to 0.199. We rounded this number to 0.2. Thus, by the strictest measure, FC values between 0.8 and 1.2 were classified as ‘neutral’. Accordingly, FC ≤0.8 unambiguously assigned a mutation as ‘deleterious’, and FC >1.2 as ‘beneficial’.

### Genome sequencing

We sequenced the genomic DNA of all biofilm populations for which we had ≥ 1 μg of DNA after extraction (6 normal and 12 gradient biofilm). For comparison, we also sequenced initial mix populations 1 and 2, 6 Liquid and 4 pellicle populations. The Illumina HiSeq2500 platform was used, with 2×125 base pairs read length. We obtained a total of 300 million reads. The reads were assembled using as reference the *B. subtilis* NCIB 3610 genome (NCBI Accession number: CP020102). Overall, 95% of all reads were successfully mapped to the reference genome with minimal coverage of x300 for all samples analyzed. The Breseq program was used to identify genomic variants, including single nucleotide polymorphisms (SNPs) and insertion-deletion polymorphisms (INDELs) ^45^ (**Data S3** & **Table S6**).

### Comparison of FC values and to GudB’s natural sequence variability

We examined whether the FC values for individual mutations, in individual conditions, and in combinations thereof, might predict whether or not a certain sequence exchange is observed, or not, amongst the sequences of naturally occurring GDHs. To this end, we constructed a number of different support vector machines (SVM) classification models with a variety of kernels (such as linear, Gaussian, polynomial etc.). The feature vector of each GudB allele was composed from the normalized FC values from specific condition. The values from replica experiments of the highly reproducible liquid conditions were averaged prior to training. Based on the multiple sequence alignment containing 1013 GDH sequences, we divided the GudB mutations in our dataset into 3 categories, which were then utilized as the prediction labels: (1) mutations seen in less than 5 natural GDH sequences (classified as ‘not present’, 66% of mutations), (2) mutations observed in 5 - 49 sequences (‘rare’, 19%) and (3) mutations present in ≥50 sequences (‘frequent’, 15%). Introducing class weights into the loss function compensated the unbalanced nature of the dataset. For each feature combination of a varied length, we built an SVM classification model and assessed its accuracy using 3-fold cross validation. Additionally, in order to reduce noise, assuming that our data belong to linear space, we extracted the first ten principal components of the feature matrix and used them as the new feature vectors for a model construction. To examine if our relatively high (>0.6) model accuracy was distributed uniformly across different classes, for each model and genotype, we recorded the predicted values during 3-fold cross-validation. Moreover, for each condition combination, and for each kernel, we built 100 different models and recorded the number of times each of the genotypes was predicted correctly.

